# Evidence for the F200Y (T<u>A</u>C) mutation conferring benzimidazole resistance in a southern USA cattle population of *Haemonchus placei* spreading from a single emergence

**DOI:** 10.1101/578922

**Authors:** Umer Chaudhry, E. M. Redman, Ray Kaplan, Thomas Yazwinski, Neil Sargison, John S. Gilleard

## Abstract

The benzimidazoles are one of the most important broad-spectrum anthelmintic drug classes for the control of parasitic nematodes of domestic animals and humans. They have been widely used in the livestock sector, particularly in small ruminants for over 40 years. This has resulted in the development and wide spread of resistance in small ruminant gastrointestinal nematode parasite species, including *Haemonchus contortus.* Recently, resistance to benzimidazole drugs has been reported in *Haemonchus placei*, but there is relatively little information on its prevalence. It is important to develop a molecular tools to identify resistance mutations in *H. placei* early in their development in order to understand the emergence and spread. Our previous study demonstrated the F200Y (T**<u>A</u>**C) mutation at their early stage in 6/9 *H. placei* populations derived from southern USA, albeit at low frequencies between 2 and 10%. The present study analysis the phylogenetics of the isotype-1 β-tubulin locus to suggest that F200Y (T**<u>A</u>**C) mutation has been spread from a single emergence in *H. placei*; likely by the anthroprogenic movement of ruminant livestock in southern USA. Population genetic data of *H. placei* using a panel of microsatellite markers revealed little genetic sub-structure, consistent with a high level of gene flow in this region. Overall, these results provide clear genetic evidence for the spread of F200Y (T**<u>A</u>**C) benzimidazoles resistance mutation to multiple different locations from a single emergence in *H. placei*.

## 1 Introduction

Gastrointestinal nematode parasites are the major cause of disease in grazing ruminants, results in billions of US dollars in production losses each year in livestock industry worldwide (Stromberg and Gasbarre, 2006). *Haemonchus placei* represents the most important and highly pathogenic parasite of large ruminants and *Haemonchus contortus* mostly infects sheep and goats, having significant economic losses in tropical and sub-tropical regions (Hoberg et al., 2004; Lichtenfels et al., 1994; Lichtenfels JR, 1994).

The mechanism of benzimidazole resistance has been studied in small ruminant nematode parasites and strong evidence exists that three different single nucleotide polymorphisms (SNPs) in the isotype-1 β-tubulin locus are responsible for resistance (Chaudhry et al., 2015a). Despite widespread global concern regarding benzimidazole resistance in *H. contortus* of small ruminants, until recently little attention has been given to the possibility of resistance develop in cattle parasites (Coles, 2002). Nonetheless, benzimidazole resistance is now emerging and represents a serious challenge to the cattle industry worldwide (Sutherland and Leathwick, 2011). There have been relatively few studies of the genetic determinants of benzimidazole resistance in *H. placei* (Ali et al., 2019a; Brasil et al., 2012).

The understanding how the benzimidazole resistance mutations emerge and spread in nematode parasite populations is an important goal. There are two ways in which adaptive mutations might develop and spread in nematode parasite populations that are under selection. First, benzimidazole resistance could emerge as a new mutation and then spread through populations by host migration, likely as a consequence of animal movements: in this case a single resistance haplotype is present in each population. This has been observed for the benzimidazole resistance conferring allele E198A (G**<u>C</u>**A) in *H. contortus* (Chaudhry et al., 2015d) and more recently F200Y (T**<u>A</u>**C) in *H. placei* (Ali et al., 2019b). Second, benzimidazole resistance could repeatedly emerge multiple time by recurrent or pre-existing mutations and migrate between nematode parasite populations as a result of host movement. In this case multiple resistance haplotypes will be present in each population. This has been proposed for benzimidazole resistance alleles F200Y (T**<u>A</u>**C) and F167Y (T**<u>A</u>**C) of *H. contortus* (Chaudhry et al., 2015a; Chaudhry et al., 2015d; Redman et al., 2015).

In the present study, we first use a panel of seven microsatellite markers to show the population genomic structure of *H. placei* and *H. contortus* from three different southern US states and then use the phylogenetic approaches to investigate the emergence and spread of benzimidazole resistance in the *H. placei* populations. The data provide clear evidence for the spread of the *H. placei* F200Y (T**<u>A</u>**C) resistance mutation from a single emergence. We also analysis the benzimidazole resistance allele frequency in seven *H. contortus* populations of sheep and goats. The F200Y (T**<u>A</u>**C) mutation was found in all seven populations at high frequency and F167Y (T**<u>A</u>**C) mutation was present in four populations relatively low frequency. The phylogenetic analysis suggests that the F200Y (TAC) and F167Y (T**<u>A</u>**C) mutations has emerge multiple independent times in the Southern USA.

## 2. Materials and Methods

### 2.1 Parasite material, genomic DNA extraction and pyrosequencing genotyping

Parasite materials were obtained from three regions of southern USA, where we anticipated a high *Haemonchus* prevalence. Adult *Haemonchus* worms were harvested from the abomasa of 10 cattle, 2 sheep and 4 goats immediately following slaughter at three different locations (Arkansas, Florida, and Georgia). Detail of the 10 cattle parasite populations has been described in a previous report (Chaudhry et al., 2015c). Two and three *Haemonchus* populations of sheep and goats, respectively, were collected from Arkansas (Pop1S, Pop2S, Pop10G, Pop11G, Pop12G) and one goat-derived *Haemonchus* population was collected from Georgia (Pop1G). Four populations (Pop2S, Pop10G, Pop11G and Pop12G) were collected directly from an abattoir, hence the host grazing history was unknown. One population (Pop1S) was collected from a farm, where sheep has been grazed on a single pasture for 6 months prior to necropsy. One population (Pop1G) was collected directly from abattoir, with no grazing history (Supplementary Table S1).

Adult worms were fixed in 80% ethanol immediately following removal from the host abomasum. The heads of individual worms were dissected and lysed in single 0.5ul tube containing 40 μl of proteinase K lysis buffer and stored at −80°C as previously described by Chaudhry et al. (2016b). 1 μl of a 1:5 dilution of a neat single worm lysate was used as PCR template and identical dilutions of lysis buffer, made in parallel, were used as negative controls. To prepare pooled lysates of each population, 1 μl aliquots of each individual neat adult worm lysate were pooled. 1 μl of a 1:50 dilution of pooled lysates was used as PCR template. Pyrosequence genotyping was performed to target the rDNA ITS-2 and codons F167Y (T**<u>A</u>**C), E198A (G**<u>C</u>**A) and F200Y (T**<u>A</u>**C) of isotype-1 β-tubulin of the *H. placei* and *H. contortus* was described in our previously studies (Chaudhry et al., 2014; Chaudhry et al., 2015b).

### 2.2 Microsatellite genotyping

Six previously published microsatellites (Hcms3561, Hcms53265, Hpms43, Hpms52, Hpms53, Hpms102) were selected as a potentially useful markers based on their properties (Chaudhry et al., 2015a; Chaudhry et al., 2016b; Santos et al., 2017). These studies were produce a clear unambiguous genotypes with either a single or double Genescan peak on single worms as anticipated for single markers in both *H. placei* and *H. contortus*. Individual worm genotyping was performed from six *H. placei* populations (Pop76C, Pop9C, Pop80C, Pop85C, Pop88C, Pop87C) and 4 *H. contortus* populations (Pop1G, Pop10G, Pop11G, Pop12G) contained F200Y (TAC) and F167Y (TAC) resistance associated SNP. A summary of primers sequences, allele ranges, PCR amplification and bioinformatics analysis was described in our previous studies (Chaudhry et al., 2016a; Santos et al., 2017)

### 2.3 Phylogenetic analysis of the isotype-1 β-tubulin locus

For the isotype-1 β-tubulin gene, a fragment zencompassing parts of exons 4 and 5 including codons F167Y (T**<u>T</u>**C-T**<u>A</u>**C), E198A (G**<u>A</u>**A-G**<u>C</u>**A) and F200Y (T**<u>T</u>**C-T**<u>A</u>**C) for *H. placei* (325bp) and H. contortus (328bp) were amplified. Pooled lysates were made of six *H. placei* populations (Pop9C, Pop76C, Pop80C, Pop87C, Pop88C, Pop85C), in which F200Y (T**<u>A</u>**C) was detected and seven *H. contortus* populations (Pop1S, Pop2S, Pop10G, Pop11G, Pop12G, Pop1G, Pop86C) in which F200Y (TAC) and F167Y (TAC) was detected. Amplicons were cloned into PJET 1.2/BLUNT vector (Thermo scientific) and sequenced using standard procedures were described by Chaudhry et al. (2015b). For the phylogenetic analysis, sequences were aligned with *H. placei* isotype-1 β-tubulin sequences (Acc No KJ598498) and edited using Geneious Pro 5.4 software (Drummond AJ, 2012). A previously described approaches were used to filter the isotype-1 β-tubulin sequences to remove SNPs occurring only once in the dataset and ensure PCR-induced mutations was not included in the analysis (Chaudhry et al., 2015d; Redman et al., 2015). The aligned sequences were then imported into the CD-HIT software (Huang et al., 2010) to calculate the number of unique haplotypes present in each population (Table 4). Construction of network tree of the isotype-1 β-tubulin haplotypes has been described in our previous studies (Chaudhry et al., 2015a; Chaudhry et al., 2016b).

## 3. Results

### 3.1 Confirmation of H. placei and H. contortus species in southern USA

Seven out of the 10 cattle populations were identified as 100% *H. placei* (P24; **<u>G</u>** genotype), one population (Pop86C from Georgia) was identified as 100% *H. contortus* (P24 **<u>A</u>** genotype), one population (Pop9C) was identified as 97% *H. contortus* (P24 **<u>A</u>** genotype) and 3% *H. placei* (P24; **<u>G</u>** genotype) and one population (H85) was identified as *H. placei* except single worm with a heterozygous **<u>A</u>**/**<u>G</u>** at position P24, suggesting that it may be *H. placei* / *H. contortus* hybrid (Supplementary Table S1 & Fig. 1). In the present study, between 29 and 32 individual *Haemonchu*s worms were pyrosequence genotyped for the rDNA ITS-2 P24 SNP (64 worms form sheep and 125 worms from goats) and all worms identified as *H. contortus* (P24 **<u>A</u>** genotype) (Supplementary Table S1 & Fig. 1).

**Fig. 1.**
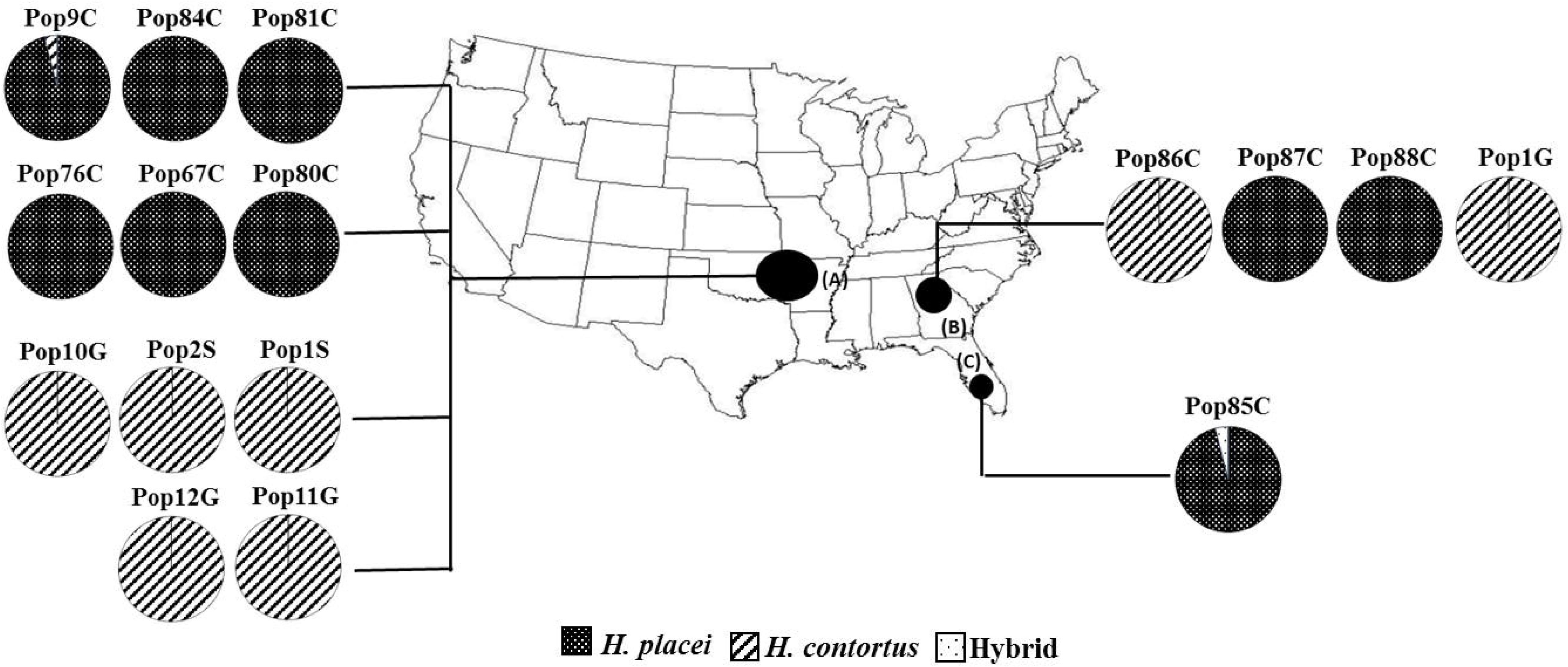
Distribution of *Haemonchus* spp. identified in southern USA. Geographic locations of abattoirs/farms are indicated with small black circles in three states (A) Arkansas (B) Georgia (C) Florida. Each pie chart represents a single parasite population taken from an individual host. The final letter of the parasite population name indicates the host species of origin (S, sheep; G, goat; C, cattle). Black shading represents worms identified as *H. placei* (Homozygous **<u>G</u>** at ITS-2 rDNA P24), vertical line shading represents worms identified as *H. contortus* (Homozygous **<u>A</u>** at ITS-2 rDNA position P24) and light dot represents worms identified as putative hybrids (heterozygous **<u>A</u>**/**<u>G</u>** at ITS-2 rDNA P24).

### 3.2 Frequency of the F167Y, E198A, F200Y polymorphisms in H. placei and H. contortus isotype-1 β-tubulin locus

Six of the 9 *H. placei* populations had F200Y (TAC) benzimidazole resistance associated SNP at frequencies between 2 to 10% (data in Supplementary Table S2). The benzimidazole resistance associated SNP F167Y (T**<u>A</u>**C) and E198A (G**<u>C</u>**A) was not detected in any of the populations. In the present study, benzimidazole resistance-associated SNPs were found in all 7 populations with the F200Y (T**<u>A</u>**C) mutation at high frequencies between 82-100% and 4 populations with the F167Y (T**<u>A</u>**C) mutation at low frequencies between 7-24% (Supplementary Table S2). The benzimidazole resistance associated SNP E198A (G**<u>C</u>**A) was not detected in any of the populations.

### 3.3 Population genetic structure of H. placei and H. conrotus

Six *H. placei* and 4 *H. contortus* populations were successfully genotyped using a panel of six microsatellite markers. To measure the level of genetic diversity between populations, the diversity index value was estimated. All populations were polymorphic at all loci, with the overall number of alleles per locus (A) ranging from 3 to 16 in *H. placei* and 2 to 10 in *H. contortus* respectively. There was a number of unique alleles (A_U_) specific to each population and broadly similar alleles were observed in each population (Table 1). There was some significant departure from Hardy-Weinberg equilibrium, even after Bonferroni correction, in 4 out of the 36 loci combinations for *H. placei* and 3 out of the 24 loci combinations for *H. contortus*, respectively (Table 1). There were no major departures from linkage equilibrium for any particular combination of loci across all populations indicating that alleles at these loci were randomly associated. *H. placei* and *H. contortus* showed a high level of overall genetic diversity in all populations, the mean allele richness (A_C_) was 7.750 ± 0.603 and 5.292 ± 0.479 respectively and expected heterozygosity (H_e_) was 0.705 (range: 0.042-0.701) and 0.488 (range: 0.048-0.546) respectively (Table 1). To measure the level of genetic structure between populations, the AMOVA and Fixation index (F_ST_) value was estimated. The percentage of variation that partitioned between 6 *H. placei* populations was 0.042% and 4 *H. contortus* populations was 0.015%. This was reflected by levels of pairwise F_ST_ estimates with a maximum of 0.09 for 13 out of 15 possible pairwise comparisons in *H. placei*, and maximum of 0.02 for 4 out of 6 possible pairwise comparisons in *H. contortus*, showing significant genetic differentiation (Table 2).

### 3.4 Haplotype distribution and the network analysis of isotype-1 β-tubulin locus of H. placei and H. conrotus

A 325bp fragment of *H. placei* isotype-1 β-tubulin was cloned and sequenced from six populations individually, DNA template was pooled from between 29 to 36 worms (Supplementary Table S1) and maximum of 12 clones were sequenced per population (Supplementary Table S3). Based on the analysis of each population separately, a single F200Y (T**<u>A</u>**C) resistance-conferring haplotype (Hr3 F200Y) was present in all six populations relatively high frequencies (Supplementary Table S3; Fig. 2A), demonstrating evidence of single emergence of benzimidazole resistance mutations. In contrast, one population (Pop87C) contained a single susceptible haplotype, three populations (Pop9C, Pop76C, Pop85C) contained a maximum of 2 susceptible haplotypes, one population (Pop87C) contained 3 susceptible haplotypes and one populations (Pop88C) contained a maximum of 4 susceptible haplotypes (Supplementary Table S3, Fig. 2A).

**Fig. 2.**
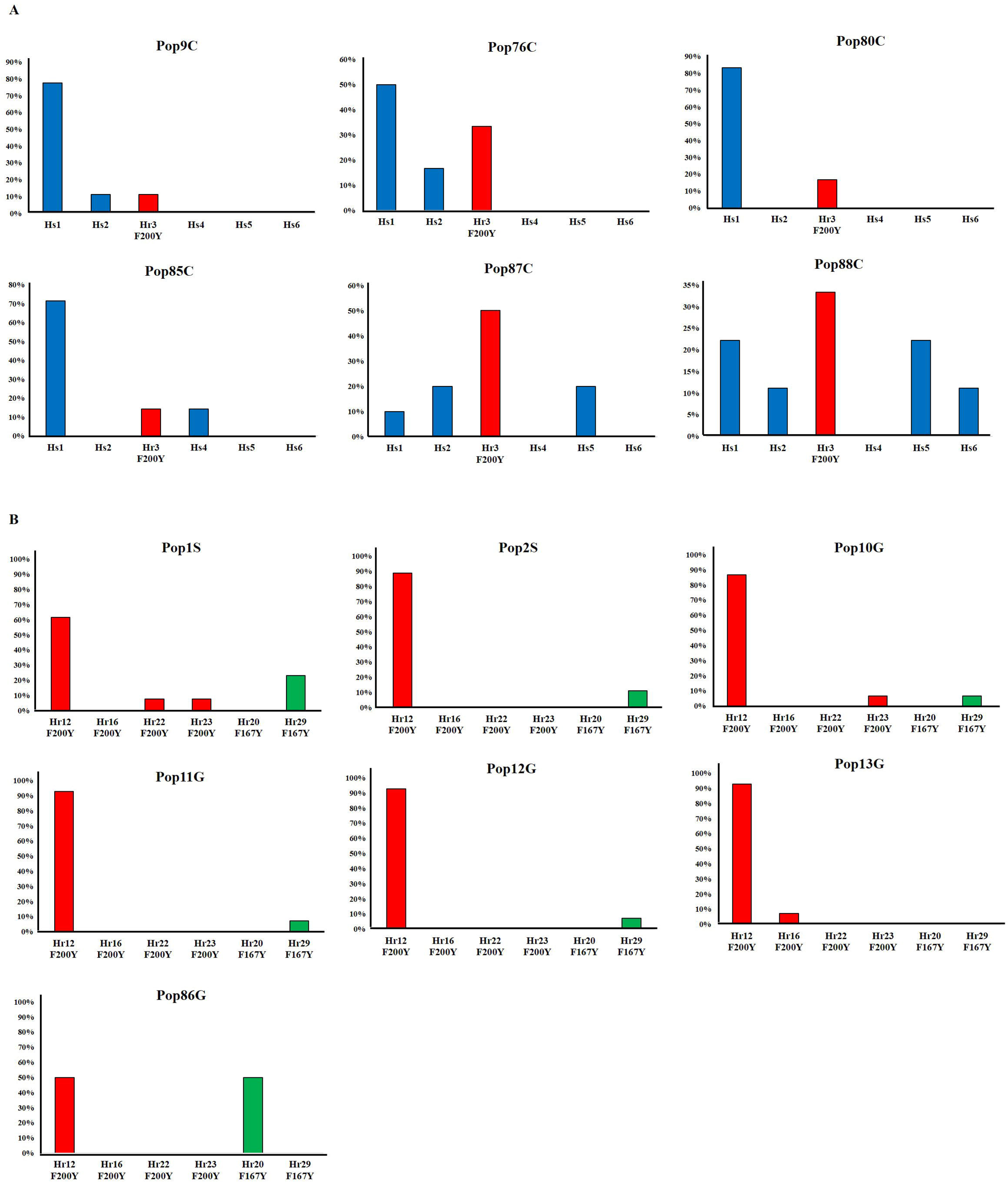
Frequency histograms showing resistant and susceptible isotype-1 β-tubulin haplotypes identified from six *H. placei* populations in panel A and seven *H. contortus* populations in (panel B). F200Y (TTC)/ F167Y (TTC)/E198A (GAA) susceptible haplotypes are shown in blue, F200Y (T**<u>A</u>**C) resistant haplotypes in red colour and F167Y (T**<u>A</u>**C) resistant haplotypes in green colour.

A total of five susceptible and one resistant unique haplotypes of *H. placei* isotype-1 β-tubulin were identified among 39 and 13 sequences of six *H. placei* populations (Supplementary Table S3). The network tree was produced to examine the phylogenetic relationship between six isotype-1 β-tubulin haplotypes (Fig. 3A). The analysis showed that the F200Y (T**<u>A</u>**C) resistance SNP was present on a single (Hr3 F200Y) haplotype in the tree (Fig. 3A). This resistance haplotype was more closely related to one or more susceptible haplotypes and represented three populations from Arkansas (Pop9C, Pop76C, Pop80C), two populations from Georgia (Pop87C, Pop88C) and one population from Florida (Pop85C) (Fig. 3A) suggests that this mutation arose once and spread to the multiple different locations of southern USA through the flow of drug resistance alleles.

**Fig. 3.**
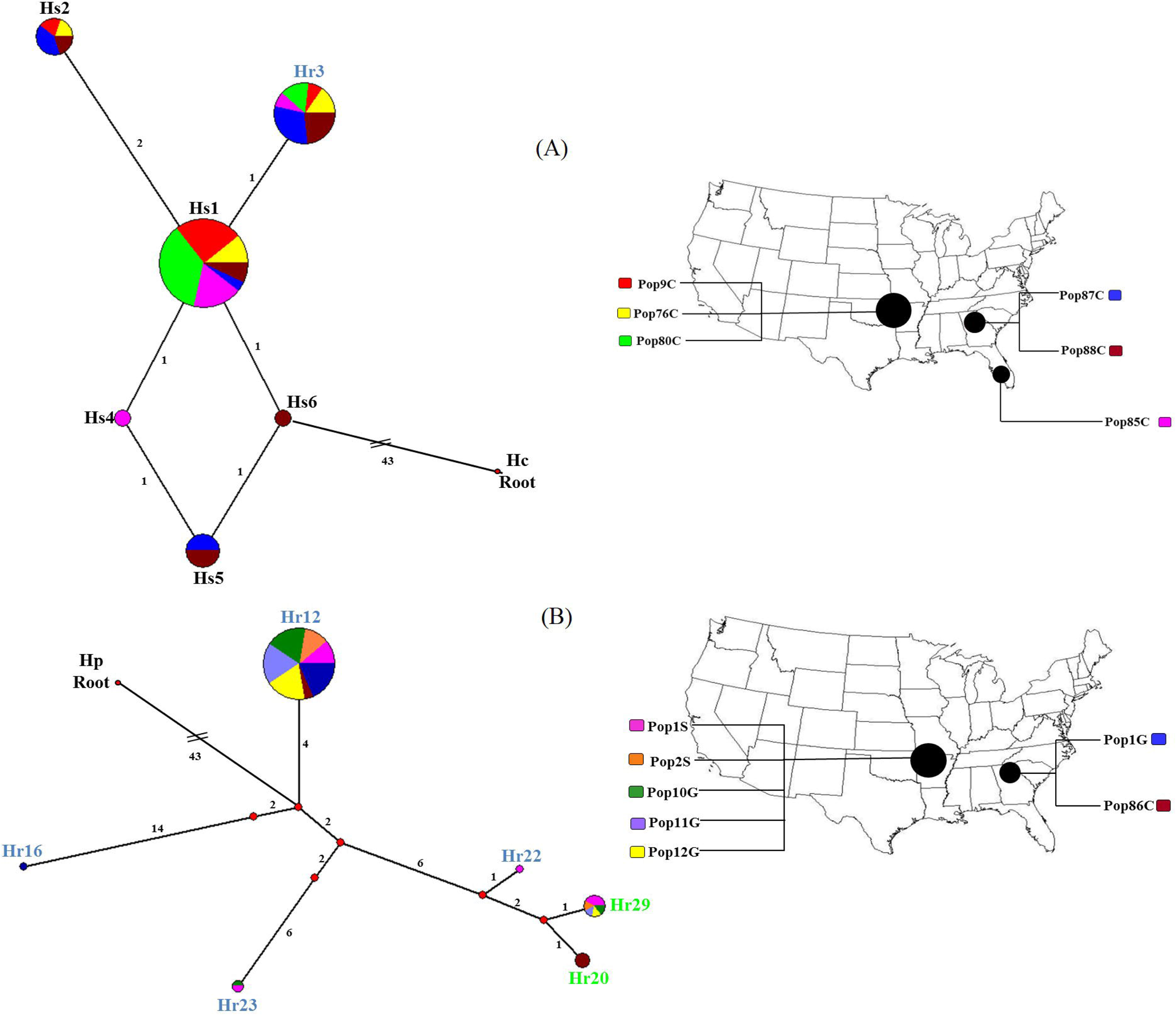
Median joining network of the *H. placei* (panel A) and *H. corturtus* (panel B) isotype-1 β-tubulin sequences generated in Network 4.6.1. A full median network containing all possible shortest trees was generated by setting the epsilon parameter equal to the greatest weighted distance (epsilon = 10). All unnecessary median vectors and links are removed with the MP option (Polzin and Daneschmand, 2003). The size of circle representing each haplotype is proportional to its frequency in the dataset and the colours in the circles reflect the spread of this haplotype in each population as indicated on the colour key on the inset map. The number of mutations separating adjacent sequence nodes or median vectors is indicated along connecting branches and the length of the lines connecting the haplotypes is proportional to the number of nucleotide changes. The most probable ancestral node is determined by rooting the network to a closely related outgroup *H. contortus* (Hc) against *H. placei* network (GenBank accession number **<u>X67489</u>**) and outgroup *H. placei* (Hp) against *H. contortus* network (GenBank accession number **<u>KJ598498</u>**). The text providing the name of each haplotype is colour coded as follows; susceptible haplotypes F200Y (T**T**C)/ F167Y (T**T**C)/E198A (G**<u>C</u>**A) is in black text; F200Y (T**A**C) resistant haplotype is in blue text; F167Y (T**A**C) resistant haplotype is in green text.

In contrast, a 328bp of *H. contortus* isotype-1 β-tubulin was cloned and sequenced from seven populations individually, DNA template was pooled from between 29 to 32 worms (Supplementary Table S1) and maximum of 15 clones were sequenced per population (Supplementary Table S3). Based on the analysis of each popualtion separately, one population (Pop1S) contained a maximum of 4 F200Y (T**<u>A</u>**C) and F167Y (T**<u>A</u>**C) resistance haplotypes, one population (Pop10G) had 3 F200Y (T**<u>A</u>**C) and F167Y (T**<u>A</u>**C) resistance haplotypes. Moreover, four populations (Pop2S, Pop11G, Pop12G, Pop86G) contained a maximum of 2 F200Y (T**<u>A</u>**C) and F167Y (T**<u>A</u>**C) resistance haplotypes and one population (Pop13G) contained a maximum of 2 F200Y (T**<u>A</u>**C) resistance haplotypes (Supplementary Table S3; Fig. 2B). In all these seven populations had high frequencies of F200Y (T**<u>A</u>**C) and F167Y (T**<u>A</u>**C) resistance-conferring haplotypes (Supplementary Table S3; Fig. 2B), demonstrating evidence of multiple time emergence of benzimidazole resistance mutations. The benzimidazole susceptible haplotypes were not detected in any of the populations.

A total of 6 resistant unique haplotypes of *H. contortus* isotype-1 β-tubulin were identified among 85 sequences of seven *H. contortus* populations (Supplementary Table S3). The network trees were produced to examine the phylogenetic relationship between six isotype-1 β-tubulin haplotypes (Fig. 3B). The analysis showed that the F200Y (T**<u>T</u>**C) resistance associated haplotype Hr12 was present in seven populations (Pop1S, Pop2S, Pop10G, Pop11G, Pop12G, Pop1G, Pop86C), Hr23 resistance haplotype present in two populations (Pop10G, Pop1S), Hr16 and Hr22 haplotype present in Pop1G and Pop1S populations from Arkansas and Georgia (Fig. 3B). Similarly, the F167Y (T**<u>T</u>**C) resistance associated haplotype (Hr29) present in five populations (Pop1S, Pop2S, Pop10G, Pop12G, Pop1G) and Hr20 resistance haplotype was present in Pop86C population from Arkansas and Georgia (Fig. 3B). The overall results suggests that F200Y (T**<u>T</u>**C) and F167Y (T**<u>T</u>**C) arose multiple time and spread to the different locations of southern USA through the flow of drug resistance alleles.

## 4. Discussion

The control of parasitic nematodes in ruminants is heavily dependent on the use of benzimidazole drug. They have been intensively used in the livestock sector for over 40 years (Chaudhry, 2015). This has led to the development of resistance in a number of small ruminant parasite species including *H. contortus* (Chaudhry, 2015). Benzimidazole resistance has been reported in closely related cattle parasite *H. placei*, but there is relatively little information on its prevalence (Ali et al., 2019a; Brasil et al., 2012). In the present study, we proposed that benzimidazole conferring resistance would be at early stage of development in *H. placei* as compared to *H. contortus* in southern USA due to less drug selection pressure in cattle as compared to small ruminants.

We demonstrated the allele frequency of known benzimidazole resistance–conferring mutations in *H. placei* populations of cattle from southern USA (Chaudhry et al., 2015c). We have detected the F200Y (T**<u>A</u>**C) mutation in 6 out of 9 *H. placei* populations. These results suggest that the F200Y (T**<u>A</u>**C) resistance mutation is likely to be present in many *H. placei* populations in southern US cattle, observed by Brasil et al. (2012) and Ali et al. (2019b). The presence of this mutation at a low frequency would not be expected to result in detectable loss of efficacy of benzimidazole drug (Ali et al., 2019a). In the present study, phylogenetic analysis of *H. placei* F200Y (T**<u>A</u>**C) resistance haplotype (Hr3) supports the hypothesis that it emerged from a single resistance mutation. Indeed, in this case, it appears that this mutation has spread to multiple different locations in southern USA and this mutation is present on a single haplotype in all 6 of the *H. placei* populations. In contrast, there was a high level of sequence diversity of susceptible haplotypes (five different susceptible haplotypes in the dataset). Hence, given this high level of susceptible haplotypes in southern USA, it would be extremely unlikely that the F200Y (T**<u>A</u>**C) mutation repeatedly arose only on the same haplotype (Hr3) in future. The fact that this mutation has single emergence illustrates the role of animal movement in the spread of this resistance allele. There have been relatively few studies of the molecular genetics of benzimidazole resistance in *H. placei* in cattle. Recently, Ali et al. (2019b) also look in to the evidence for the single emergence of the F200Y (T**<u>A</u>**C) resistance mutation and spread in Pakistani *H. placei* populations through unregulated animal movement.

We have confirmed the allele frequency of benzimidazole resistance mutations in *H. contortus* populations of sheep and goats. The F200Y (T**<u>A</u>**C) mutation was present in all 7 populations at high frequency and F167Y (T**<u>A</u>**C) mutation was found in 4 out 7 populations relatively low frequency. The F200Y (T**<u>A</u>**C) resistance conferring SNP is widespread and repeatedly present at high frequencies in many developed countries including New Zealand, Sweden, France, Australia and UK (Bisset SA et al., 2014; Hoglund et al., 2009; Kotze et al., 2012; Redman et al., 2015; Silvestre and Humbert, 2002). In contrast, the other known benzimidazole resistance mutation F167Y (T**<u>A</u>**C) is more variable in its prevalence. Although the F167Y (T**<u>A</u>**C) has been detected in France and Canada, it is generally less widespread and at much lower frequencies than F200Y (T**<u>A</u>**C) mutation except in UK (Barrère et al., 2012; Barrere et al., 2013a; Barrere et al., 2013b; Silvestre and Cabaret, 2002). In the present work, we also found that benzimidazole resistance are well advance in southern USA, and the F200Y (T**<u>A</u>**C) is the predominant resistance mutation in *H. contortus* populations.

There have studies investigating the emergence and spread of benzimidazole resistance in *H. contortus* populations of small ruminants, where resistance is at early stages. These studies demonstrated that F200Y (T**<u>A</u>**C) and F167Y (T**<u>A</u>**C) mutation emerged multiple time in small ruminants (Chaudhry et al., 2016b; Redman et al., 2015). Knowledge of the emergence of *H. contortus* resistance in those studies makes it very likely that the movement of livestock is an important element in the spread of benzimidazole resistance alleles in small ruminants. However, this is difficult to investigate and demonstrate in the present southern USA, where benzimidazole resistance is well advanced almost at the level of fixation, for example F200Y (T**<u>A</u>**C) mutation is present in all populations examined at high frequencies. Therefore, the diversity of the resistance alleles and the complexity of their relationships to susceptible alleles, makes it difficult to use genetic analysis to definitively demonstrate that particular resistance allele has emerge and then spread from one location to another in southern USA.

The data reported in the present study reveal a high genetic diversity among *H. placei* populations (allele richness 7.750 ± 0.603, expected heterozygosity 0.705) and *H. contortus* (allele richness 5.292 ± 0.47, expected heterozygosity 0.488). Similar studies have described a evidence of high genetic diversity in *H. contortus* populations (Chaudhry et al., 2015a; Chaudhry et al., 2016b; Hunt et al., 2008; Redman et al., 2015), but there are very few reports on *H. placei* (Ali et al., 2018). The impact of the high level of genetic diversity will not influence the benzimidazole resistance mutation rate under the impact of drug selection pressure, when compared both susceptible and resistant *H. contortus* and *H. placei* populations (Ali et al., 2018; Chaudhry et al., 2016a). In the present study, a low, but significant level of genetic differentiation has also been observed in the populations of *H. placei* (F_st_ estimates maximum of 0.09) and *H. contortus* (F_st_ estimates maximum of 0.02). Similar finding suggest that genetic differentiation does occur at low but significant level in *H. contortus* populations (Chaudhry et al., 2015a; Chaudhry et al., 2016b; Hunt et al., 2008; Redman et al., 2015) and a few reports in *H. placei* (Ali et al., 2018). The consequences of low levels of population genetic differentiation in part reflect high levels of animal movement (Hunt et al., 2008). If the gene flow is high, the F200Y (T**<u>A</u>**C), E198A (G**<u>C</u>**A) and F167Y (T**<u>A</u>**C) resistance mutations potentially spread in the regions under the influence of drug selection. These observations have also been recognized in *H. contortus* and *H. placei* studies (Ali et al., 2019b; Chaudhry et al., 2015a; Chaudhry et al., 2016b; Redman et al., 2015; Yin et al., 2016).

## Conclusions

The present study in the southern USA, where benzimidazole resistance in *H. placei* is at early stages, provides a simpler situation from where to draw conclusions as compared to *H. contortus*. The single emergence of F200Y (T**<u>A</u>**C) resistance mutations in *H. placei* and its subsequent spread is clearly defined. The way in which this mutation has become widespread provides a clear illustration of the role of migration in the spread of resistance alleles. The results of the high level of genetic diversity in southern US *H. placei*, may be explained by large effective population size and high mutation rate. Furthermore, low levels of genetic differentiation results into the animal movement may have led to the increase of gene flow of *H. placei* populations in this region. There is need for the better understanding of farming practices and the parasite epidemiology to inform sustainable control of gastrointestinal nematode parasites in different part of the world. This includes better emphasis on the importance of biosecurity measures and quarantine dosing in managing the emergence of resistance in the regions where there is considerable movement of animals. Our results suggest that the spread of resistance alleles in different locations play an important role in producing the complex patterns of resistance seen at the early stage of development.

## Acknowledgements

We would like to thank Natural Sciences and Engineering Research Council of Canada (NSERC) for funding support (Grant number RGPIN/371529-2209) as well as the NSERC-CREATE Host Pathogen Interactions (HPI) graduate training program at the University of Calgary. Work at the Roslin Institute uses facilities funded by the Biotechnology and Biological Sciences Research Council (BBSRC).

