## Supplementary material for "Evidence for the F200Y (T<u>A</u>C) mutation conferring benzimidazole resistance in a southern USA cattle population of *Haemonchus placei* spreading from a single emergence": Table 1

**Table 1:** Population genetic data for each microsatellite marker from six *H. placei* and four *H. contortus* populations based on panel of 6 microsatellite loci.

| ***H. placei*** | | | | | | | |
| --- | --- | --- | --- | --- | --- | --- | --- |
| *Population* | *Hcms43 (8^b^)* | *Hcms52 (21^b^)* | *Hcms53 (7^b^)* | *Hcms102 (14^b^)* | *Hcms3561 (9^b^)* | *Hcms5365 (7^b^)* | *All loci* |
| *Pop76C (28^a^)* |  |  |  |  |  |  |  |
| N_o_ | 0 | 1 | 0 | 0 | 0 | 0 |  |
| A_U_ | 7(0) | 15(1) | 6(0) | 11(1) | 6(0) | 3(0) | 49(2) |
| p-value | 0.0385 | 0.0477 | 0.3314 | 0.5550 | 0.0158 | 1.0000 |  |
| H_e_ | 0.8103 | 0.9070 | 0.7896 | 0.8837 | 0.6662 | 0.1370 | 0.6990 |
| H_o_ | 0.7500 | 0.8148 | 0.8214 | 0.7857 | 0.3928 | 0.1428 | 0.6179 |
| *Pop9C (22^a^)* |  |  |  |  |  |  |  |
| N_o_ | 0 | 2 | 0 | 0 | 0 | 0 |  |
| A_U_ | 8(0) | 15(0) | 5(0) | 8(1) | 7(0) | 4(1) | 47(2) |
| p-value | 0.9528 | 0.1182 | 0.8642 | 0.4742 | 0.1571 | 1.0000 |  |
| H_e_ | 0.7547 | 0.9346 | 0.7526 | 0.7896 | 0.7029 | 0.3583 | 0.7154 |
| H_o_ | 0.7727 | 0.9000 | 0.7727 | 0.7272 | 0.5454 | 0.4090 | 0.6878 |
| *Pop80C (26^a^)* |  |  |  |  |  |  |  |
| N_o_ | 0 | 0 | 0 | 0 | 0 | 0 |  |
| A_U_ | 7(0) | 13(1) | 4(0) | 10(0) | 4(0) | 3(0) | 41(1) |
| p-value | 0.5974 | 0.3985 | 0.3354 | 0.3262 | 0.0050 | 1.0000 |  |
| H_e_ | 0.8137 | 0.9012 | 0.7096 | 0.8506 | 0.6289 | 0.3054 | 0.7016 |
| H_o_ | 0.7692 | 0.9230 | 0.6153 | 0.7692 | 0.4230 | 0.3461 | 0.6410 |
| *Pop85C (31^a^)* |  |  |  |  |  |  |  |
| N_o_ | 0 | 0 | 0 | 0 | 0 | 1 |  |
| A_U_ | 7(0) | 16(0) | 8(1) | 8(1) | 6(0) | 3(0) | 48 (2) |
| p-value | 0.9687 | 0.3154 | 0.5429 | 0.3471 | 0.0076 | 0.1999 |  |
| H_e_ | 0.7667 | 0.9360 | 0.7403 | 0.6266 | 0.6663 | 0.3406 | 0.6794 |
| H_o_ | 0.7096 | 0.9354 | 0.7741 | 0.8387 | 0.4838 | 0.2666 | 0.6681 |
| *Pop88C (27^a^)* |  |  |  |  |  |  |  |
| N_o_ | 0 | 1 | 0 | 0 | 0 | 0 |  |
| A_U_ | 8(0) | 16(0) | 7(0) | 7(1) | 7(0) | 5(1) | 50(2) |
| p-value | 0.2362 | 0.6290 | 0.9341 | 0.1066 | 0.0013 | 1.0000 |  |
| H_e_ | 0.7232 | 0.9027 | 0.7526 | 0.3053 | 0.7302 | 0.1775 | 0.5986 |
| H_o_ | 0.8148 | 0.8846 | 0.7777 | 0.2222 | 0.4074 | 0.1851 | 0.5486 |
| *Pop87C (30^a^)* |  |  |  |  |  |  |  |
| N_o_ | 0 | 0 | 2 | 1 | 0 | 0 |  |
| A_U_ | 6(0) | 13(0) | 4(0) | 10(0) | 4(0) | 5(1) | 42(1) |
| p-value | 0.4166 | 0.3893 | 0.2677 | 0.0608 | 0.0000 | 1.0000 |  |
| H_e_ | 0.7565 | 0.8977 | 0.7337 | 0.8663 | 0.6655 | 0.2552 | 0.6958 |
| H_o_ | 0.7333 | 0.9666 | 0.6785 | 0.6896 | 0.3333 | 0.2758 | 0.6129 |
| ***H. contortus*** | | | | | | | |
| *Population* | *Hcms43 (4^b^)* | *Hcms52 (14^b^)* | *Hcms53 (5^b^)* | *Hcms102 (6^b^)* | *Hcms3561 (5^b^)* | *Hcms5365 (11^b^)* | *All loci* |
| *Pop1G (26^a^)* |  |  |  |  |  |  |  |
| N_o_ | 0 | 0 | 0 | 8 | 0 | 0 |  |
| A_U_ | 2(1) | 9(2) | 5(0) | 2(1) | 3(0) | 6(0) | 27(4) |
| p-value | 1.0000 | 0.0179 | 0.2264 | 1.0000 | 0.2083 | 0.17225 |  |
| H_e_ | 0.0384 | 0.8106 | 0.6078 | 0.1079 | 0.4954 | 0.6040 | 0.4440 |
| H_o_ | 0.0384 | 0.5600 | 0.5769 | 0.1111 | 0.3846 | 0.4800 | 0.3585 |
| *Pop10G (26^a^)* |  |  |  |  |  |  |  |
| N_o_ | 0 | 0 | 0 | 0 | 0 | 0 |  |
| A_U_ | 3(0) | 5(0) | 5(0) | 4(1) | 5(2) | 9(1) | 31(4) |
| p-value | 1.0000 | 0.1938 | 0.2735 | 0.1738 | 0.0562 | 0.1743 |  |
| H_e_ | 0.1802 | 0.6757 | 0.5935 | 0.2149 | 0.4208 | 0.5799 | 0.4441 |
| H_o_ | 0.1923 | 0.6923 | 0.5000 | 0.1923 | 0.3461 | 0.5000 | 0.4038 |
| *Pop11G (28^a^)* |  |  |  |  |  |  |  |
| N_o_ | 0 | 0 | 0 | 0 | 0 | 0 |  |
| A_U_ | 3(0) | 10(2) | 5(0) | 3(0) | 3(0) | 8(0) | 32(2) |
| p-value | 1.000 | 0.6746 | 0.7149 | 0.8480 | 0.0960 | 0.1587 |  |
| H_e_ | 0.1370 | 0.8097 | 0.6039 | 0.4694 | 0.4642 | 0.7961 | 0.5467 |
| H_o_ | 0.1428 | 0.8214 | 0.6428 | 0.4285 | 0.3571 | 0.8214 | 0.5357 |
| *Pop12G (29^a^)* |  |  |  |  |  |  |  |
| N_o_ | 0 | 0 | 0 | 0 | 1 | 0 |  |
| A_U_ | 3(0) | 8(0) | 5(0) | 4(0) | 3(0) | 7(1) | 30(1) |
| p-value | 1.0000 | 0.2343 | 0.9197 | 0.6122 | 0.0734 | 0.3337 |  |
| H_e_ | 0.2266 | 0.6824 | 0.6474 | 0.3720 | 0.3221 | 0.5850 | 0.4726 |
| H_o_ | 0.2500 | 0.5714 | 0.6428 | 0.3928 | 0.2222 | 0.4642 | 0.4239 |

N_o_, apparent null homozygotes, i.e. number of worms in the population which failed to give an amplification product for a particular marker; A, number of alleles; A_U_, number of alleles unique to that population; H_e_, expected heterozygosity; H_o_, observed heterozygosity; P-values indicate a significant deviation from Hardy–Weinberg equilibrium following bonferroni correction.

^a^ Total number of individuals genotyped for each population is given in parenthesis under the population name.

^b^ Total number of alleles for each marker across all populations is given in parenthesis below each marker name.

.
