## Supplementary material for "Evidence for the F200Y (T<u>A</u>C) mutation conferring benzimidazole resistance in a southern USA cattle population of *Haemonchus placei* spreading from a single emergence": Table 1

**Table 2.** Pairwise F_ST_ comparison between the six *H. placei* and four *H. contortus* populations from three different geographical locations in Southern USA using six distinct microsatellite markers.

***H. placei	H. contortus***

|  | Pop76C | Pop9C | Pop80C | Pop85C | Pop88C |
| --- | --- | --- | --- | --- | --- |
| Pop9C | **0.02089** |  |  |  |  |
| Pop80C | 0.00735 | **0.01853** |  |  |  |
| Pop85C | **0.05024** | 0.00284 | **0.04111** |  |  |
| Pop88C | **0.09582** | **0.06766** | **0.09652** | **0.04451** |  |
| Pop87C | **0.02994** | **0.02087** | **0.02781** | **0.02669** | **0.06586** |

|  | Pop1G | Pop10G | Pop11G |
| --- | --- | --- | --- |
| Pop10G | **0.01716** |  |  |
| Pop11G | **0.02304** | 0.01082 |  |
| Pop12G | **0.02529** | 0.00273 | **0.01612** |
