## Supplementary Table S1 for "Evidence for the F200Y (T<u>A</u>C) mutation conferring benzimidazole resistance in a southern USA cattle population of *Haemonchus placei* spreading from a single emergence"

**Supplementary Table S1.** rDNA ITS-2 species-specific SNP (P24) genotypes identified by pyrosequencing of the worms from southern USA. Populations of *Haemonchus* collected from cattle, sheep and goats host from Arkansas, Florida and Georgia region.

| **US field populations** | **Host** | **No. of *H. contortus* worms** | **No. of *H. placei*** **worms** | **No. of *Heterozygous*** **worms** | **Worm number** | **Origin** | **Location** |
| --- | --- | --- | --- | --- | --- | --- | --- |
| Pop9C | Cattle | 1 | 31 |  | 32 | Abattoir | Arkansas |
| Pop67C | Cattle |  | 32 |  | 32 | Abattoir | Arkansas |
| Pop76C | Cattle |  | 29 |  | 29 | Abattoir | Arkansas |
| Pop80C | Cattle |  | 32 |  | 32 | Abattoir | Arkansas |
| Pop81C | Cattle |  | 32 |  | 32 | Abattoir | Arkansas |
| Pop84C | Cattle |  | 32 |  | 32 | Abattoir | Arkansas |
| Pop85C | Cattle |  | 36 | 1 | 37 | Abattoir | Florida |
| Pop87C | Cattle |  | 32 |  | 32 | Abattoir | Georgia |
| Pop88C | Cattle |  | 32 |  | 32 | Abattoir | Georgia |
| Pop86C | Cattle | 29 |  |  | 29 | Abattoir | Georgia |
| Pop1S | Sheep | 32 |  |  | 32 | Farm | Arkansas |
| Pop2S | Sheep | 32 |  |  | 32 | Abattoir | Arkansas |
| Pop10G | Goat | 32 |  |  | 32 | Abattoir | Arkansas |
| Pop11G | Goat | 29 |  |  | 29 | Abattoir | Arkansas |
| Pop12G | Goat | 32 |  |  | 32 | Abattoir | Arkansas |
| Pop1G | Goat | 32 |  |  | 32 | Abattoir | Georgia |
