## Supplementary Table S2 for "Evidence for the F200Y (T<u>A</u>C) mutation conferring benzimidazole resistance in a southern USA cattle population of *Haemonchus placei* spreading from a single emergence"

| **US field populations** | **Host** | **No of worms in each pool** | **F167Y** | | **E198A** | | | **F200Y** | | **Location** |
| --- | --- | --- | --- | --- | --- | --- | --- | --- | --- | --- |
|  |  |  | TTC | T**A**C | GAA | G**C**A | TTC | | T**A**C |  |
| *H. placei* |  |  |  |  |  |  |  | |  |  |
| Pop9C | Cattle | 31 | 100% | 0% | 100% | 0% | 98% | | 2% | Arkansas |
| Pop67C | Cattle | 32 | 100% | 0% | 100% | 0% | 100% | | 0% | Arkansas |
| Pop76C | Cattle | 29 | 100% | 0% | 100% | 0% | 98% | | 2% | Arkansas |
| Pop80C | Cattle | 32 | 100% | 0% | 100% | 0% | 98% | | 2% | Arkansas |
| Pop81C | Cattle | 32 | 100% | 0% | 100% | 0% | 100% | | 0% | Arkansas |
| Pop84C | Cattle | 32 | 100% | 0% | 100% | 0% | 100% | | 0% | Arkansas |
| Pop85C | Cattle | 36 | 100% | 0% | 100% | 0% | 96% | | 4% | Florida |
| Pop87C | Cattle | 32 | 100% | 0% | 100% | 0% | 90% | | 10% | Georgia |
| Pop88C | Cattle | 32 | 100% | 0% | 100% | 0% | 98% | | 2% | Georgia |
| *H. contortus* |  |  |  |  |  |  |  | |  |  |
| Pop86C | Cattle | 29 | 92% | 8% | 100% | 0% | 5% | | 95% | Georgia |
| Pop1S | Sheep | 32 | 100% | 0% | 100% | 0% | 0% | | 100% | Arkansas |
| Pop2S | Sheep | 32 | 76% | 24% | 100% | 0% | 18% | | 82% | Arkansas |
| Pop10G | Goat | 32 | 94% | 8% | 100% | 0% | 10% | | 90% | Arkansas |
| Pop11G | Goat | 29 | 93% | 7% | 100% | 0% | 8% | | 94% | Arkansas |
| Pop12G | Goat | 32 | 100% | 0% | 100% | 0% | 0% | | 100% | Arkansas |
| Pop1G | Goat | 32 | 100% | 0% | 100% | 0% | 0% | | 100% | Georgia |

**Supplementary Table S2.** Allele frequency (%) of SNPs that resulted in an amino acid change at codons F167Y (TTC/TAC), E198A (GAA/GCA) and F200Y (TTC/TAC) in isotype-1 β-tubulin obtained from nine *H. placei* and seven *H. contortus* populations from southern USA.
