## Supplementary Table S3 for "Evidence for the F200Y (T<u>A</u>C) mutation conferring benzimidazole resistance in a southern USA cattle population of *Haemonchus placei* spreading from a single emergence"

**Supplementary Table S3.** Total number of sequences, resistant and susceptible isotype-1 β-tubulin haplotypes identified from six *H. placei* and seven *H. contortus* populations*.*

| ***H. placei* populations** | **No. of Sequences** | **F200Y (TAC) resistant haplotypes (%)** | **Susceptible haplotypes (%)** | **Total no of haplotypes** |
| --- | --- | --- | --- | --- |
| Pop9C | 9 | 1 (11.11%) | 2 (88.81%) | 3 |
| Pop76C | 6 | 1 (33.33%) | 2 (66.60%) | 3 |
| Pop80C | 12 | 1 (16.66%) | 1 (83.33%) | 2 |
| Pop85C | 7 | 1 (14.20%) | 2 (85.62%) | 3 |
| Pop87C | 10 | 1 (50%) | 3 (50%) | 4 |
| Pop88C | 9 | 1 (33.33%) | 4 (66.66%) | 5 |
| R* | 13 | 1 (25%) |  |  |
| S* | 39 |  | 5 (75%) |  |
| Total | 52 | 1 | 5 | 6 |

| ***H. contortus* populations** | **No. of Sequences** | **F200Y (TAC) resistant haplotypes (%)** | **F167Y (TAC) resistant haplotypes**  **(%)** | **Susceptible haplotypes**  **(%)** | **Total no of haplotypes** |
| --- | --- | --- | --- | --- | --- |
| Pop1S | 13 | 3 (76.73%) | 1 (23.07%) | 0 (0%) | 4 |
| Pop2S | 9 | 1 (88.88%) | 1 (11.11%) | 0 (0%) | 2 |
| Pop10G | 15 | 2 (93.26%) | 1 (6.60%) | 0 (0%) | 3 |
| Pop11G | 14 | 1 (92.80%) | 1 (7.10%) | 0 (0%) | 2 |
| Pop12G | 14 | 1 (92.80%) | 1 (7.10%) | 0 (0%) | 2 |
| Pop13G | 14 | 2 (100%) | 0 (0%) | 0 (0%) | 2 |
| Pop86C | 6 | 1 (50%) | 1 (50%) | 0 (0%) | 2 |
| R* | 85 | 4 (88.16%) | 2 (12.40%) |  |  |
| S* | 0 |  |  | 0 (0%) |  |
| Total | 85 | 4 | 2 |  | 6 |

* R & S indicated overall resistant and susceptible sequences and haplotypes in six *H. placei* and seven *H. contortus* populations.
